# Correlating glycoforms of DC-SIGN with stability using a combination of enzymatic digestion and ion mobility MS

**DOI:** 10.1101/2020.04.22.055046

**Authors:** Hsin-Yung Yen, Idlir Liko, Joseph Gault, Di Wu, Weston B. Struwe, Carol V. Robinson

## Abstract

The immune scavenger protein DC-SIGN interacts with glycosylated proteins and has a putative role in facilitating viral infection. How these recognition events take place with different viruses is not clear and the effects of glycosylation on the folding and stability of DC-SIGN have not been reported. Here, we develop and apply a mass spectrometry-based approach to both uncover and characterise the effects of O-glycans on the stability of DC-SIGN. We first quantify the Core 1 & 2 O-glycan structures on the carbohydrate recognition and extracellular domains of the protein via sequential exoglycosidase sequencing. We then use ion mobility mass spectrometry to show how specific O-glycans, and/or single monosaccharide substitutions, alter both the overall collision cross section and the gas-phase stability of the glycoprotein isoforms of DC-SIGN. We find that rather than the mass or length of glycoprotein modifications, the stability of DC-SIGN is better correlated with the number of glycosylation sites. Collectively, our results exemplify a combined multi-dimensional MS approach, proficient in evaluating protein stability in response to both glycoprotein macro- and micro-heterogeneity and adding structural detail to the infection enhancer DC-SIGN.

---

While much current interest focuses on the glycosylation status of the viral spike protein of SARS-CoV-2, few studies have addressed the role of receptor glycosylation. DC-SIGN (the innate immune receptor dendritic cell-specific intercellular adhesion molecule-3 grabbing non-integrin) has been implicated as an ‘infection enhancer’ in previous reports of coronavirus epidemics^[1,2]^. This property is attributed to the ability of DC-SIGN either to recognize self- or other pathogenic carbohydrates. As such DC-SIGN is proposed to play an unfavourable role in coronavirus infections, enhancing circulation of virions through multivalent interactions of high-mannose type viral glycans via its carbohydrate recognition domain (CRD)^[3]^. This ability to increase circulation of viral particles has prompted efforts to develop glycomimetic drugs for DC-SIGN, as CRD antagonists, to inhibit host-virus interactions and infection^[4]^. To the best of our knowledge however, DC-SIGN, a C-type lectin expressed on the surface of dendritic cells and macrophages, has yet to be reported as O-glycosylated and the extent and heterogeneity of these glycoforms has yet to be defined.

Here we use DC-SIGN as a challenging test case to develop and apply high-resolution native MS and ion mobility (IM) instrumentation to study the effects of glycosylation on the biologically relevant forms of the receptor. DC-SIGN is a type II membrane protein comprising three main domains: a cytoplasmic region, a transmembrane segment, and an extracellular domain (ECD). Although DC-SIGN is known to be tetrameric, enabling multivalent interaction with pathogens, a complete structure is not available; largely due to the intrinsic flexibility of the ECD. The ECD can be divided into two distinct regions: a neck region involved in tetramerization of the receptor and CRD, which mediates the molecular recognition processes. A model of the ECD has been proposed, based on small angle X-ray scattering data^[5]^.

Our principal goal is to understand the impact of glycosylation on the stability of the DC-SIGN receptor by developing and applying a combined native MS approach to assign the overall glycan occupancy (macroheterogeneity) and to characterize its detailed structural information (microheterogeneity). These two types of data are generally obtained separately through orthogonal MS and liquid chromatography techniques. Here, we show that both macro- and microheterogeneity information can be acquired simultaneously through a single MS experiment using specific monosaccharide glycosidases in sequence. We further show, using IM and collision induced unfolding (CIU) measurements, that specific glycan structures, as well as the extent of their occupancy, affect the stability of the intact glycoprotein. Intriguingly we find that the overall mass of the glycan is less important in affecting the stability of DC-SIGN than the number of glycosylation sites. Taken together this approach therefore provides a means to gain both structural and biophysical information for this intact folded glycoprotein that is not accessible by other static or ensemble-based methods.

We began our investigation by expressing and purifying DC-SIGN CRD from human embryonic kidney (293T) cells. The native MS spectrum of this protein revealed two major charge states (8^+^ and 9^+^) with seven clear proteoforms within each distribution (**Figure 1a**). The theoretical mass of non-glycosylated DC-SIGN CRD is 19128 Da. The lowest mass observed was 21014.5 ± 0.4 Da with additional peaks indicating the presence of further PTMs. The mass differences between these peaks correspond to monosaccharides with distinct numbers of hexose (+162 Da), N-acetylhexosamine (+203 Da) and N-acetylneuraminic acid (+292 Da) residues. The difference of +1886.5 ± 0.4 Da is consistent with 2 hexoses (Hex), 2 N-acetylhexosamines (HexNAc) and 4 N-acetylneuraminic acid residues (Neu5Ac is referred to as sialic acid herein). Further mass shifts on the CRD domain are from Hex-HexNAc disaccharide additions plus a further 2 sialic acids. At this stage of the analysis HexNAc monosaccharides are unspecified (blue/yellow squares) as they can be either N-acetylgalactosamine and/or N-acetylglucosamine residues. We then recorded the native mass spectrum of the full ECD domain and observed a closely similar glycosylation pattern (**Figure 1b**). We conclude that the CRD/ECD glycan modifications range between (Hex-HexNAc)_2-6_ with either 4 or 6 sialic acids - the most glycosylated forms having 6 Hex, 6 HexNAc and 6 Neu5Ac monosaccharides (**Figure 2a**).

**Figure 1.**
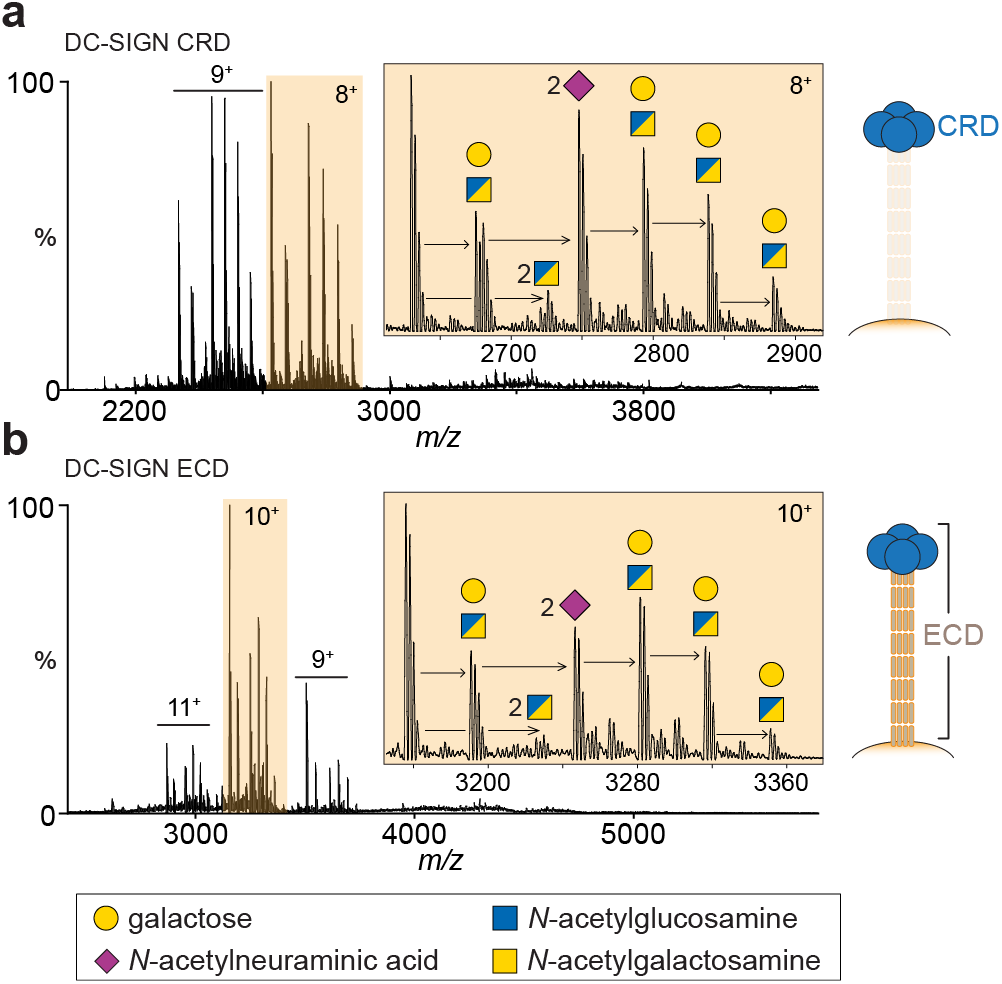
Identification of O-glycosylation on DC-SIGN. (**a**) Native mass spectrum of the DC-SIGN carbohydrate recognition domain (CRD) monomer displaying two major charge states (8^+^ and 9^+^). Peaks corresponding to glycoforms with mass shifts of hexose (shown as galactose), *N*-acetylhexosamine and *N*-acetylneuraminic acid (sialic acid) residues are shown (8^+^ charge state). A split blue/yellow box denotes a residue that can be either *N*-acetylglucosamine or *N*-acetylgalactosamine. (**b**) Native mass spectrum of the extracellular domain (ECD) monomer (10+ charge state), containing the CRD plus the neck domain, reveals a similar glycosylation pattern.

**Figure 2.**
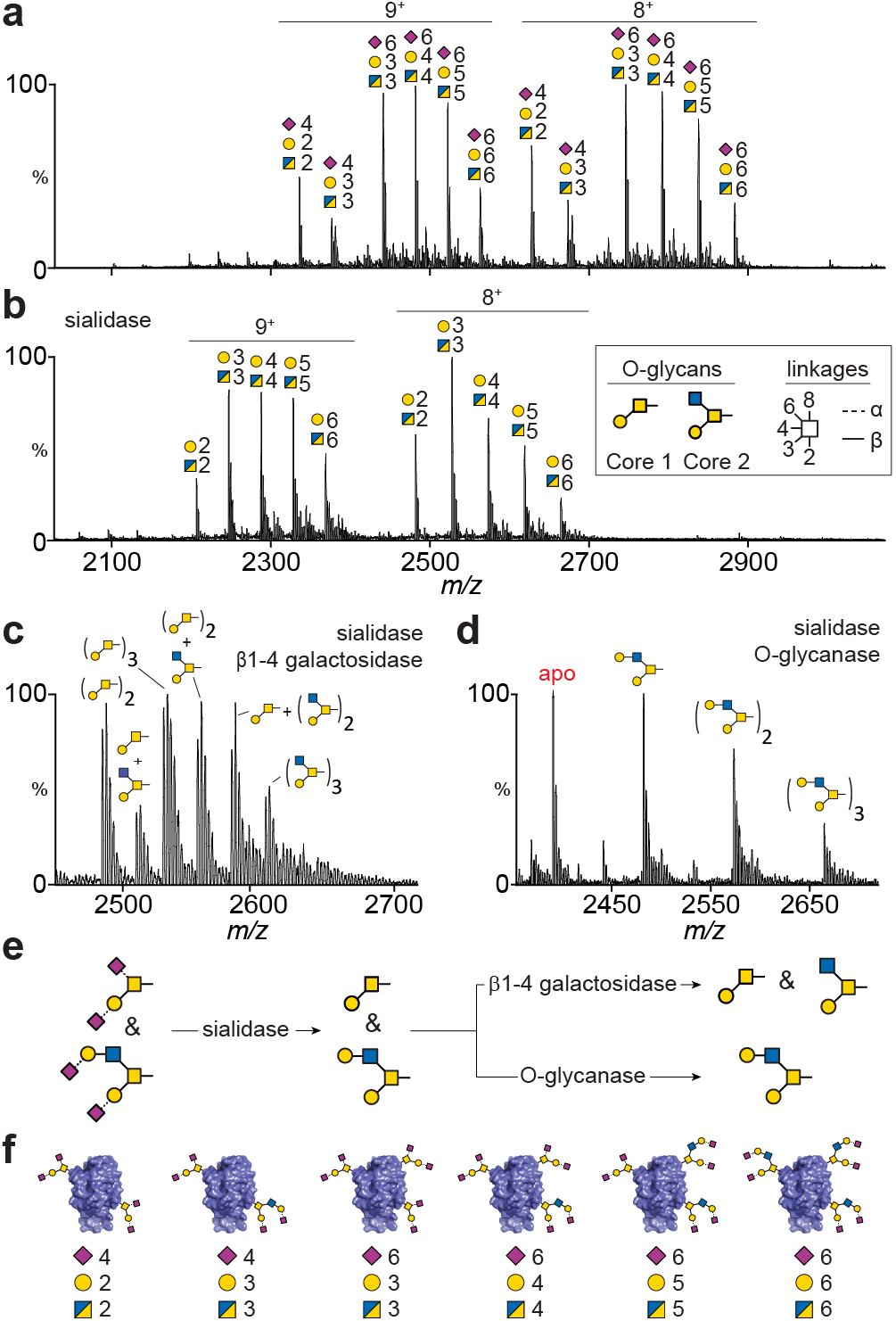
Exo- and endoglycosidase digestions of DC-SIGN CRD. (**a**) Mass spectrum of CRD without enzymatic treatment indicates extensive sialylation of at least six glycoforms in each charge state. (**b**) Sialidase digestion results in five peaks with different hexose and *N*-acetylhexosamine compositions. (**c**) Sequential digestion with sialidase and β1-4 galactosidase reveals presence of both Core 1 and Core 2 O-glycan structures, (inset in (**b**)). (**d**) Replacing β1-4 galactosidase with O-glycanase, which is specific for Core 1 O-glycans, results in formation of the non-glycosylated CRD (apo peak) and identification of β1-4 galactose extensions on Core 2 O-glycans. (**e**) Schematic of CRD glycan structures generated from exo- and endoglycosidase digestions. (**f**) DC-SIGN CRD glycan macroheterogeneity and microheterogeneity identified from exoglycosidase sequencing.

Since the CRD of DC-SIGN lacks an Asn-x-Ser/Thr sequon, the oligosaccharide PTMs we detected are unlikely to be N-linked glycans. The numbers of monosaccharides observed by MS are however characteristic of O-linked glycans, which typically are smaller than N-glycans, and occur on Ser/Thr amino acids. Covalent attachment of N-acetylgalactosamine is the first step in O-linked glycan biosynthesis, followed by addition of galactose, N-acetylglucosamine and sialic acid residues. We cannot infer any structural or occupancy information from observations made in **Figure 1** as these values could arise from glycoforms comprised of six Core 1 O-glycans or any number of extended O-glycan structures.

To overcome this mass degeneracy and probe these glycan structures/microheterogeneity in more detail, we used a limited number of exo- and endoglycosidase enzymatic digestions specific for the type and linkage of individual monosaccharides. Exoglycosidases are commonly used in glycomic analyses for trimming individual residues sequentially from the non-reducing end of oligosaccharides, typically once detached and purified from the underlying protein backbone. Glycan structures (microheterogeneity) become apparent by combining known enzyme specificity with observed peak shifts. Exoglycosidase sequencing has tremendous potential for native MS of glycoproteins, by uncovering macroheterogeneity and microheterogeneity information simultaneously.

We applied this combined enzymatic MS approach to DC-SIGN CRD first using neuraminidase (sialidase) to remove α2-3,6,8 linked sialic acids from DC-SIGN CRD. From the six species identified in **Figure 2a**, removal of sialic acids now resulted in five glycoforms, with (Hex-HexNAc)_2-6_ compositions (**Figure 2b**). Combining sialidase treatment with a second enzyme (β1-4 galactosidase) we observed removal of β1-4 linked galactoses yielding six major glycoforms (**Figure 2c**) with considerable structural information emerging. The lowermost *m/z* peak is CRD with two Core 1 structures (Galβ1-3GalNAc-Ser/Thr; yellow circle and square, **Figure 2, inset**). We are confident in this assignment as other O-glycan Cores or extended O-glycan structures can be excluded because they would have an odd value of HexNAc residues, such as an extended Core 3 (**Supplemental Figure 1**). The next peak is 203 Da greater in mass and can be assigned to CRD consisting of one Core 1 and one Core 2 (GlcNAcβ1-6(Galβ1-3)GalNAc-Ser/Thr) glycan. The remaining four peaks in **Figure 2c** are CRD glycoforms with (Core 1)_3_, (Core 1)_2_(Core 2)_1_, (Core 1)_1_(Core 2)_2_ and (Core 2)_3_ structures. From this spectrum we can conclude that CRD has between two to three Core 1 and Core 2 O-linked glycans.

With this data alone, the assignment of Core 2 glycans cannot be definitive as these compositions are also equivalent to a Core 3 structure (GlcNAcβ1-3GalNAc-Ser/Thr). Core 3 and Core 1 O-glycans are susceptible to O-glycanase, so CRDs with Core 1 and Core 3 glycans would result in apo (non-glycosylated) forms which were detected (**Figure 2d**). To confirm our assignment (i.e. the absence of Core 3 structures), we treated the CRD with O-glycanase (following sialidase digestion) to remove unsubstituted Core 1 structures. This resulted in a fraction of apo CRD, which would have previously had two or three Core 1 O-glycans, with the remaining CRD glycoforms consisting of 1 to 3 extended Core 2 structures (**Figure 2d**). This result confirms our glycan assignments in **Figure 2c**. Each Core 2 glycan structure contains a single galactose residue (yellow circle), linked β1-4 to the upper N-acetylglucoasmine residue (blue square). The β1-4 galactosidase and O-glycanase digestions (**Figure 2e**) therefore confirm both the number of attached O-glycans (2 to 3) and their specific structures; Core 1 and β1-4 galactose extended Core 2 with one or two sialic acids linked to penultimate galactoses (**Figure 2f**).

Although the presence of O-glycans on DC-SIGN has yet to be reported, there is considerable evidence supporting their occurrence. NetOGlyc, a neural network learning algorithm trained by proteome-wide discovery of O-glycosyaltion sites, predicted DC-SIGN to have three possible O-glycan sites (Ser 383, Ser393 and Thr398)^[6]^. These sites were also identified using IsoGlyP (http://isoglyp.utep.edu), which allows for selection of specific polypeptide GalNAc transferases (ppGalNAc Ts) in the prediction calculation. Of the 20 known GalNAc Ts, which have different properties that govern peptide O-glycosylation, HEK293 cells used in this study express GalNAc T1, T2, T3 and T11 (included in our IsoGlyP search). The equivalent search of the DC-SIGN ECD domain did not change the location or number of O-glycosylation predictions.

Next, we used ion mobility mass spectrometry (IM-MS) to probe the stability and structure of CRD glycoforms. In addition to mass, IM-MS reports the rotationally average collision cross section (CCS) of proteins, which relates to their overall size and shape, and consequently can be used to evaluate changes in 3-dimensional structure. We focused on sialidase/O-glycanase treated CRD (herein denoted as CRD^S/O^) due to its homogeneous glycosylation (i.e. all glycans on CRD^S/O^ are galactose-extended Core 2 structures). Besides, the presence of non-glycosylated proteoforms, this analysis provided a benchmark for interpreting CCS glycoprotein data. This is based on prediction of IM-MS derived CCS values from crystal structures, however, to our knowledge only non-glycosylated proteins have been considered in this way.

Recording the arrival time distributions (ATDs) we observed peak maxima at 4.1 ms (non-glycosylated), 4.5 ms (1 O-glycan), 4.9 ms (2 O-glycans) and 5.2 ms (3 O-glycans) (**Figure 3a & 3b).**The non-glycosylated CRD^S/O^ CCS (1570 Å^2^) matched the theoretical value (1563 Å^2^) calculated from the CRD crystal structure without glycosylation (**Figure 3d**)^[7]^. CCS measurements of CRD^S/O^ with increasing numbers of glycans (1-3) revealed progressively larger structures (**Figure 3c**). With the addition of each O-glycan, the CCS increased in size by ~62 ± 2 Å^2^ from the non-glycosylated CRD^S/O^. Changes in glycoprotein CCSs were therefore consistent and glycan-specific, pointing to the potential for IM to explore gas-phase structural variations of glycoproteins.

**Figure 3.**
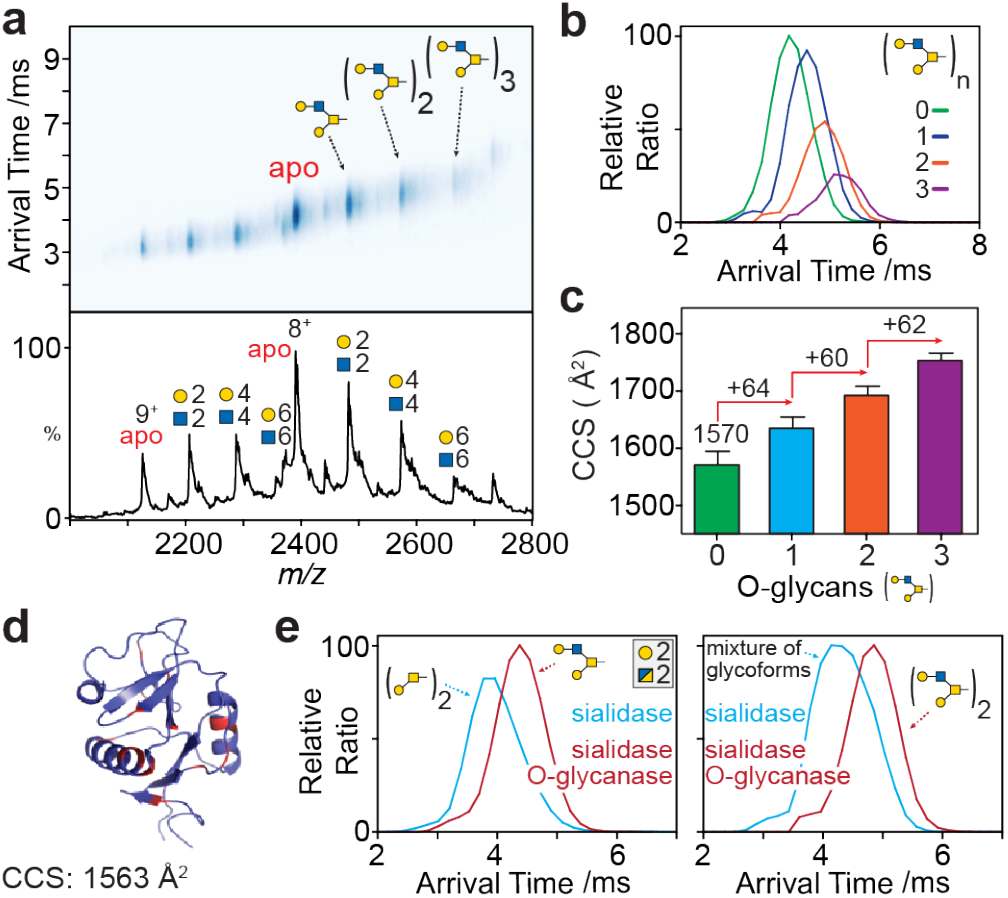
Ion mobility mass spectrometry of CRD glycoforms. (**a**) drift-plot and MS spectrum of CRD after sialidase and O-glycanase digestions (CRD^S/O^). Extracted arrival time distributions (**b**) and collision cross section values (**c**) of CRD^S/O^ (8^+^ charge state). (**d**) Crystal structure of non-glycosylated DC-SIGN CRD (PDB: 1SL4) with potential O-glycosylation sites (Ser/Thr residues) highlighted in red. (**e**) ATDs of CRD glycoforms (Hex-HexNAc)2 of identical mass; one with two Core 1 glycans (blue trace) and one with a single Core 1 glycan (red).

To explore further the ability of IM to probe these structural variations we compared CRD of similar mass but different glycosylation. The *m/z* of CRD (Gal-GalNAc)^2^ after sialidase digestion (termed CDR^S^) is the same as CRD^S/O^ with a single galactose-extended Core 2 O-glycan. As discussed above, the CDR^S^ (Gal-GalNAc)_2_ glycoform has two Core 1 structures, meaning two glycan sites are occupied, as opposed to the single extended Core 2 glycan on CRD^S/O^. The IM-MS arrival times of these two glycoforms were noticeably different (~17%) revealing distinct CCSs (**Figure 3e**). This suggests Core 1 structures induce glycoprotein compaction (i.e. lower drift time, 3.8 ms) compared to the single extended Core 2 glycoform (4.1 ms), despite its decrease in mass.

CRDs with two additional monosaccharides (Hex-HexNAc)_4_ were subsequently examined (**Figure 3e, right**). For CRD^S^, a broad ATD was observed, attributed to the presence of multiple glycoforms. By contrast, the ATD for CRD^S/0^, which has two Core 2 glycans, was narrow and more symmetric, implying fewer structures and consistent with a single proteoform. Furthermore, the ATDs increased with the addition of only 2 monosaccharides (from 3.8 to 4.1 ms for CDRS and 4.5 to 4.9 ms for CRD^S/0^) further highlighting the contribution of glycans to gas-phase protein structures. Considering the minor contribution of carbohydrate to the overall mass of the glycoprotein, which is 3.6% for CRDs with (Hex-HexNAc)_2_ and 7.1% with (Hex-HexNAc)_4_ monosaccharides, the difference in ATDs among isomeric glycoforms is apparent. The notion that sugars collapse fully onto the amino acid backbone and play only nominal roles in gas-phase structures is therefore inconsistent with our observations.

To assess how these changes in glycosylation affect protein stability we employed a protein unfolding approach using IM to follow CIU, induced by elastic collisions with a neutral gas in the mass spectrometer^[8,9]^. During a CIU experiment, the collision voltage is increased incrementally, and the protein undergoes unfolding through transition states of different CCSs. The contribution of glycosylation to protein stability can then be compared, similarly as previously described with lipid binding to membrane proteins^[10]^. Here, CIU was used to explore stability effects due to glycan microheterogeneity (neutral vs. charged), glycan macroheterogeneity as well as isobaric glycoforms (i.e. CRDs with different carbohydrate compositions but equivalent masses).

We examined the CIU profile of four CRD glycoforms, generated by glycosidase digestion, (**Figure 4a**). A single well-defined transition was observed for each CIU experiment (**Figure 4b**). We noted however that the collision voltage at which glycoform unfolding occurred varied depending on the glycan composition and occupancy. Non-glycosylated CRDs (*m/z* 2128, 19152 Da) unfolded at the lowest voltage (20 V) while CRDs with a single Core 2 O-glycan (m/z 2210, 19890 Da) unfolded at 23 V. The CRD with the greatest mass (*m/z* 2339, 21051 Da) assigned to two glycans (Hex_2_HexNAc_2_Neu5Ac_4_) had a similar unfolding pattern with a transition at 28 V. The equivalent CRD without sialic acid (2 Core 1 glycans; *m/z* 2210, 19890 Da) required significantly higher voltages to induce unfolding (37 V). In summary the greatest resistance to unfolding (stability) was observed for a CRD with two neutral O-glycans, followed by a glycoform with two negatively charged glycans followed by a CRD with a single O-glycan which was more stable than the apo protein (**Figure 4c**).

**Figure 4.**
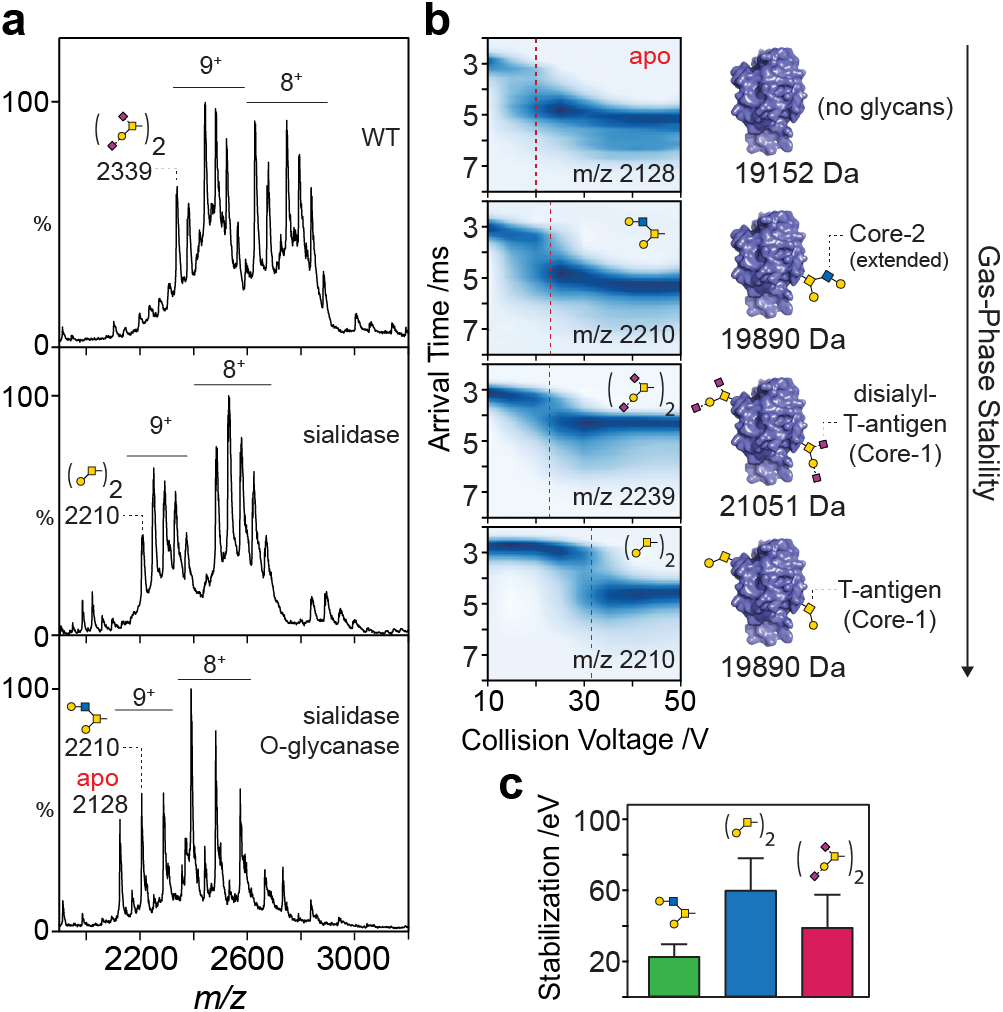
Gas-phase unfolding of DC-SIGN CRD glycoforms. (**a**) Native MS spectra of CRD untreated (top), sialidase digested (middle) and sialidase and O-glycanase digested (bottom). The four glycoforms used for unfolding experiments are labelled in each spectrum. (**b**) Corresponding collision induced unfolding plots with unfolding transitions (dotted red line) of four CRD glycoforms with varying degrees of glycosylation (right) shown at the right. (**c**) Calculated stability of CRDs with one or two O-glycans ± sialic acid residues.

These results point to variations in biophysical properties arising from protein glycosylation and a possible role for sialic acids (negatively charged monosaccharide residues) in reducing stability and/or arrangements of glycoproteins in the gas-phase. Interestingly our results indicate that the number of glycans the contributing factor to glycoform stability over the length of glycans, in agreement with a previous in silico predictions^[11]^. Together these results highlight the potential of native IM-MS to investigate the stabilization effect of individual glycans which is challenging, if not unfeasible, using other methodologies.

Given the vast diversity of glycosylation patterns evident for eukaryotic proteins, it is likely that the effects of individual glycans will be highly specific to each individual protein. Nonetheless, native mass spectrometry has great potential to explore the correlations between stability and glycosylation for any protein, as well as uncovering micro- and macroheterogenity information in a single experiment. Combining ion mobility and exoglycosidases, of which are available, means analyses can be tailored accordingly. Here, we showed how this approach not only identified O-glycosylation of DC-SIGN but enable us to characterise the extent and effects of glycosylation. These results will lead to a greater understanding of its self-recognition and potential roles in enhancing viral infections, which in turn inform therapeutic targeting.

## Abbreviations

ATD: arrival time distribution
CCS: collision cross section
CIU: collision induced unfolding
CRD: carbohydrate recognition domain
DC-SIGN: dendritic cell-specific intercellular adhesion molecule-3 grabbing non-integrin
ECD: extracellular domain
IM-MS: ion mobility mass spectrometry

## Acknowledgments

CVR is funded by a Wellcome Trust Investigator Award (104633/Z/14/Z), an ERC Advanced Grant ENABLE (641317) and an MRC Programme Grant (MR/N020413/1). J Gault is a Junior Research Fellow at The Queen’s College, Oxford. We thank Niclas Karlsson, University of Gothenburg, for discussions.

## Author contributions

Conceptualization: H.Y., I.L., W.B.S. and C.V.R.; Investigation: H.Y., I.L., D.W. and W.B.S.; Formal Analysis: H.Y.; Visualization: H.Y. and W.B.S.; Writing – Original Draft: H.Y. and W.B.S.; Writing – Review & Editing: all authors

## Experimental Section

### Construct design and protein purification

The genes of DC-SIGN extracellular domain (ECD; amino residue 66-404) and carbohydrate recognition domain (CRD; amino residue 250-404) were subcloned to pHLsec vector for protein overexpression in HEK293T cells. The condition medium harvested from the cells 48 hours post transfection by polyethylenimine, branched average Mw ~25,000Da was harvested for IMAC purification. Sample was loaded into a Histrap column pre-equilibrated with binding buffer (20 mM Tris-HCl, pH7.4, 150mM NaCl and 20 mM imidazole) at flow rate 1ml/min, and protein was eluted with the same buffer containing 500 mM imidazole in a step elution method. The protein fractions were combined for buffer exchange in 20 mM Tris-HCl, pH7.4, 150mM NaCl to remove imidazole and protein was snap frozen and stored at −80°C. For glycosidase treatment, purified recombinant protein (0.5 ug/ul) was incubated at 37°C overnight with O-glycosidase, β1-4 galatosidase or α2-3,6,8 Neuraminidase (New England BioLabs) at concentration of 5000, 1 and 12 units/ul, respectively.

### Non-denatured mass spectrometry for DC-SIGN

DC-SGIN CRD was buffer exchanged into 200 mM ammonium acetate, and immediately introduced into a modified Q-Exactive mass spectrometer (prototype Q-Exactive EMR) (Thermo Fisher) according to a previously reported method^[12]^. Overall a low voltage gradient was applied to transfer optics prior to trapping ions ions in the higher-energy collisional dissociation (HCD) cell. A low HCD activation voltage (15V) was used for protein desolvation, in order to avoid protein unfolding and fragmentation of glycan post-translational modifications. For analysis of DC-SIGN ECD, protein sample was pre-treated with 1% acetic acid to dissociate protein complexes prior introducing into mass spectrometer. Spectra were acquired with five microscans and averaged with a noise level parameter of 3. Pressure in the HCD cell was increased (measured using UHV pressure ~1.05 × 10-9 mbar) to allow better trapping and transmission of protein ions. Data was analysed by using Xcalibur 2.2 SP1.48.

### Ion mobility analysis for DC-SIGN

The collisional cross section (CCS) of DC-SIGN was measured using a modified SynaptG2-Si high definition mass spectrometer. Parameters used for analysis were the following: capillary, cone, trap and transfer collision energy were set at 1.3 kV, 50 V, 10 V and 7 V, respectively. The backing pressure was set at 6–8 mbar, and the pressure in the drift cell was set at 4.7 × 10−1 bar. The wave velocity and wave height for IMS cell is 500 m/s and 13 V whereas 248 m/s and 8V for Transfer cell. The drift time of four standard proteins (bovine serum albumin, pyruvate kinase, alcohol dehydrogenase, concanavalin A) were acquired under the same instrumental parameters for CCS calculation of DC-SIGN by a home-made software PULSAR^[13]^. Theoretical CCSs of DC-SIGN were calculated using the projection approximation method implemented in MOBCAL and scaled using a scaling factor of 1.14^[14]^. For protein unfolding experiments, the drift time for each charge state of protein was obtained under a collisional energy ramp in CID cell with 5V intervals to determine the unfolding pathway of DC-SIGN. The initial position of each unfolding species was assigned, and its intensity was extracted across all collisional voltages to generate the unfolding model by PULSAR. The stabilization effect (eV) was calculated by the sum of differences of midpoint voltage for each unfolding species in comparison to the non-glycosylated apo CRD, multiplying by individual protein charge state in order to account for charge-dependent factor in protein unfolding.

