## Supplementary Information for "Correlating glycoforms of DC-SIGN with stability using a combination of enzymatic digestion and ion mobility MS"

### Supplementary Material

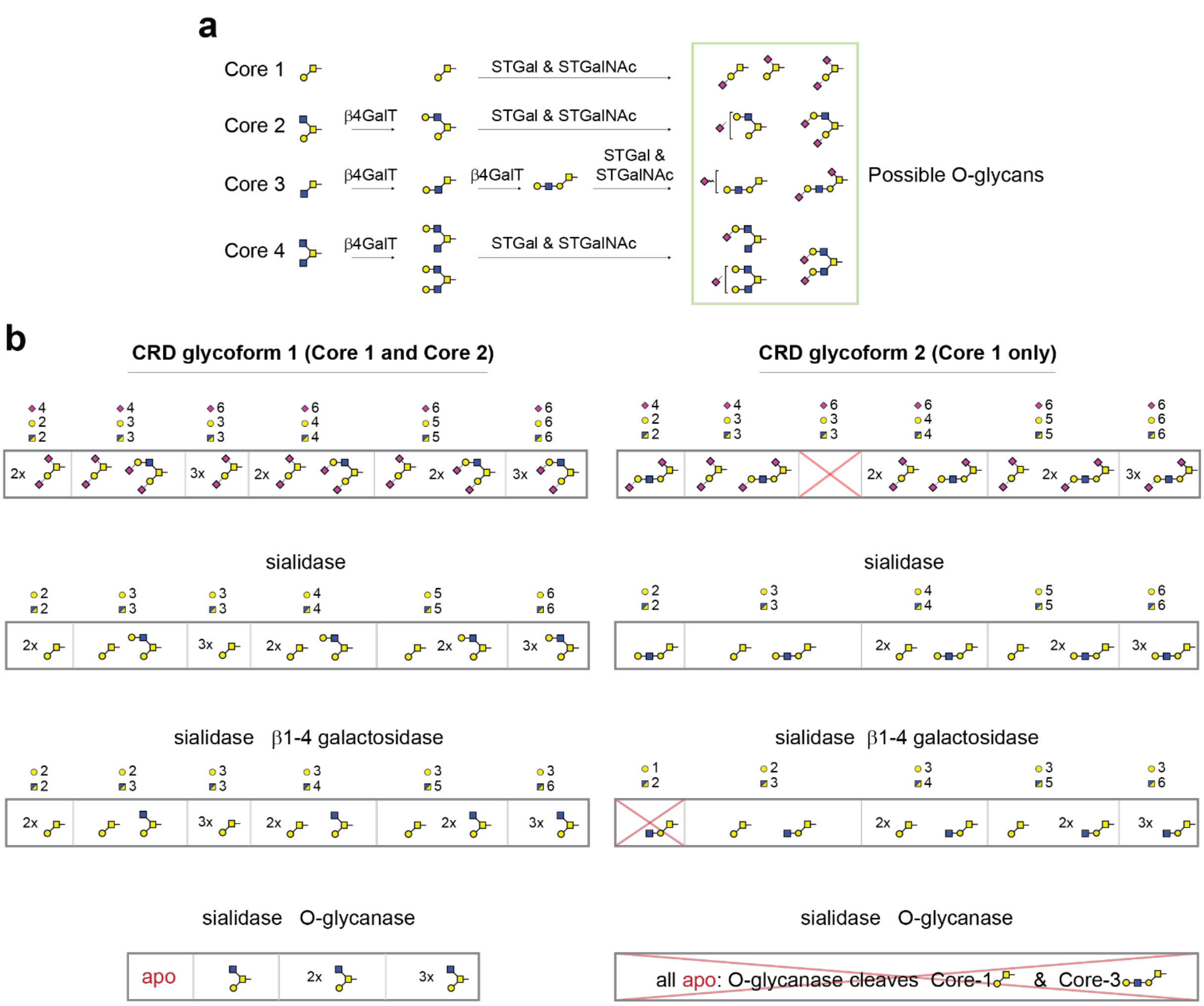


**Supplementary Figure 1: Theoretical O-glycans on DC-SIGN CRD.** (**a**) Possible O-glycans assembled from Core 1,2,3,4 structures based on addition of β1-4 galactose and α2-3,6 *N*-acetylneuraminic acid. (**b**) Corresponding to exoglycosidase digestions of theoretical O-glycans in (a) show DC-SIGN CRD contain both Core 1 and Core 2 structures.

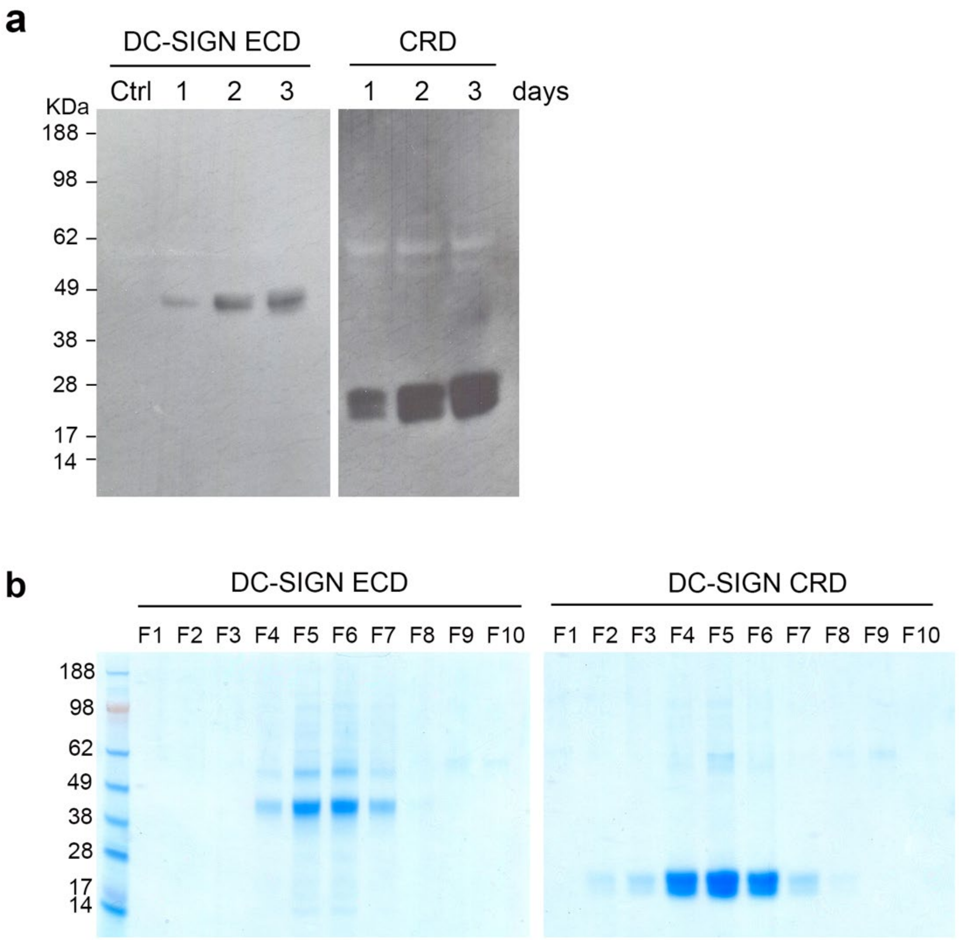


**Supplementary Figure 2: Purification of DC-SIGN CRD and ECD.** (**a**) Western blotting of condition medium harvested at different time point post transfection by polyethylenimine. (**b**) SDS-PAGE of purified DC-SIGN CRD and ECD from the elution fractions of IMAC.
